# Completing the ENCODE3 compendium yields accurate imputations across a variety of assays and human biosamples

**DOI:** 10.1101/533273

**Authors:** Jacob Schreiber, Jeffrey Bilmes, William Stafford Noble

**Affiliations:** Paul G. Allen School of Computer Science and Engineering, University of Washington; Department of Electrical Engineering, University of Washington; Department of Genome Sciences, University of Washington

## Abstract

**Motivation:** Recent efforts to describe the human epigenome have yielded thousands of uniformly processed epigenomic and transcriptomic data sets. These data sets characterize a rich variety of biological activity in hundreds of human cell lines and tissues (“biosamples”). Understanding these data sets, and specifically how they differ across biosamples, can help explain many cellular mechanisms, particularly those driving development and disease. However, due primarily to cost, the total number of assays that can be performed is limited. Previously described imputation approaches, such as Avocado, have sought to overcome this limitation by predicting genome-wide epigenomics experiments using learned associations among available epigenomic data sets. However, these previous imputations have focused primarily on measurements of histone modification and chromatin accessibility, despite other biological activity being crucially important.

**Results:** We applied Avocado to a data set of 3,814 tracks of data derived from the ENCODE compendium, spanning 400 human biosamples and 84 assays. The resulting imputations cover measurements of chromatin accessibility, histone modification, transcription, and protein binding. We demonstrate the quality of these imputations by comprehensively evaluating the model’s predictions and by showing significant improvements in protein binding performance compared to the top models in an ENCODE-DREAM challenge. Additionally, we show that the Avocado model allows for efficient addition of new assays and biosamples to a pre-trained model, achieving high accuracy at predicting protein binding, even with only a single track of training data.

**Availability:** Tutorials and source code are available under an Apache 2.0 license at https://github.com/jmschrei/avocado.

## 1 Introduction

Recently, several scientific consortia have sought to collect large sets of genomic, transcriptomic and epigenomic data. For example, since its inception in 2003, the NIH ENCODE Consortium [1] has generated over 10,000 human transcriptomic and epigenomic experiments. Similar efforts include Roadmap Epigenomics [2], modENCODE [3], the International Human Epigenome Consortium [4], mouseENCODE [5], PsychENCODE [6], and GTEx [7]. These projects have varied motivations, but all spring from the common belief that the generation of massive and diverse high-throughput sequencing data sets can yield valuable insights into molecular biology and disease.

Unfortunately, the resulting data sets are usually incomplete. In the case of ENCODE, this incompleteness is by design. Faced with a huge range of potential cell lines and primary cell types to study (referred to hereafter using the ENCODE terminology “biosample”), ENCODE investigators made the strategic decision to perform “tiered” analyses. Thus, some “Tier 1” biosamples were analyzed using a large number of different types of sequencing assays, whereas biosamples assigned to lower tiers were analyzed in less depth. This strategy allowed ENCODE to cover many biosamples while also allowing researchers to examine a few biosamples in great detail. In other cases, even for a consortium such as GTEx, which aims to systematically characterize a common set of tissue types across a set of individuals using a fixed set of assays, missing data is unavoidable due to the cost of sequencing and loss of samples during processing. Given the vast space of potential biosamples to study and the fact that new types of assays are always being developed to characterize new phenomena, the sparsity of these compendia is likely to increase over time.

This incompleteness can be problematic. For example, many large-scale analysis methods have trouble handling missing data. Despite the benefit that additional measurements may offer, many analysis methods discard assays that have not been systematically been performed in the biosamples of interest. More critically, many biomedical scientists want to exploit these massive, publicly funded consortium data sets but find that the particular biosample type that they study was relegated to a lower tier and hence is only sparsely characterized.

Imputation methods address this problem by filling in the missing data with computationally predicted values. Imputation is feasible in part due to the structured nature of consortium-style data sets, in which data from high-throughput sequencing experiments can be arranged systematically along axes such as “biosample” and “assay.” The first epigenomic imputation method, ChromImpute [8], trains a separate machine learning model for each missing experiment, deriving input features from the same row or column in the data matrix, i.e., training from experiments that involve the same biosample but a different assay or the same assay but a different biosample. A second method, PREDICTD [9], takes a more wholistic approach, first organizing the entire data set into a 3D tensor (assay × biosample × genomic position) and then training an ensemble of machine learning models that each jointly decompose all experiments in the tensor into three matrices, one for each dimension. PREDICTD imputes missing values by linearly combining values from these three matrices. Most recently, a third method, Avocado [10], extends PREDICTD by replacing the linear combination with a non-linear, deep neural network, and by modeling the genomic axis at multiple scales, thereby achieving significantly more accurate imputations without the need to train an ensemble of models.

All three of these existing imputation methods rely upon a common data set. In creating ChromImpute, Ernst and Kellis assembled what was, at the time, one of the largest collections of uniformly processed epigenomic and transcriptomic data, derived from 1,122 experiments from the Roadmap Epigenomics and ENCODE consortia. To allow for direct comparison between methods, both PREDICTD and Avocado relied upon a subset of 1,014 of those experiments. Since 2015, however, the amount of available data has increased tremendously. Here, we report the training of Avocado on a data set derived from the ENCODE compendium that contains 3,814 tracks from 400 biosamples and 84 assays (Figure 1). This ENCODE2018-Core data set is 3.4 times larger than the original ChromImpute data set. We demonstrate that this increase in size leads to a concomitant improvement in predictive accuracy.

**Figure 1:**
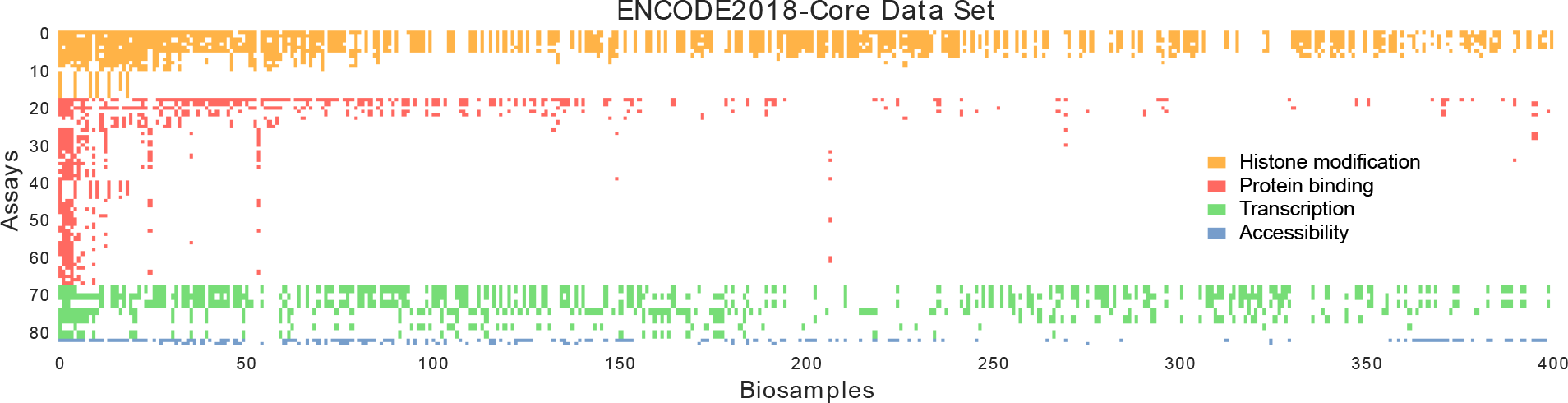
The ENCODE2018-Core data matrix. In the matrix, columns represent biosamples and rows represent assays. Colors correspond to general types of assays (histone modification ChiP-seq in orange, transcription factor ChIP-seq in red, RNA-seq in green, and chromatin accessibility in blue). Biosamples are sorted by the total number of assays performed in them, and assays are first grouped by their type before being sorted by the number of biosamples that they have been performed in.

Furthermore, whereas the ChromImpute data set included only chromatin accessibility, histone modification, and RNA-seq data, the ENCODE2018-Core data set also includes ChIP-seq measurements of the binding of transcription factors (TF) and other proteins, such as CTCF and POLR2A (referred to hereafter, for simplicity, as “transcription factors,” despite the differences in their biological roles). Accurate prediction of TF binding in a cell type-specific fashion is an extremely challenging and well-studied problem (reviewed in [11]). We demonstrate that, by leveraging the large and diverse ENCODE2018-Core data set, Avocado achieves high accuracy in prediction of TF binding, outperforming several state-of-the-art methods.

Finally, we demonstrate a practically important feature of the Avocado model, namely, that the model can be easily extended to apply to newly or very sparsely characterized biosamples and assays via a simple transfer learning approach. Specifically, we demonstrate how a new biosample or assay can be added to a pre-trained Avocado model by fixing all of the existing model parameters and only training the new assay or biosample factors. We do this using experiments from a second dataset, ENCODE2018-Sparse, that contains 3,056 experiments from biosamples that are sparsely characterized and from assays that have been performed in only few biosamples. We find that the model can yield high quality imputations for transcription factors that are added in this manner, and that these imputations can outperform the ENCODE-DREAM challenge participants even when trained using a single track of data. Finally, we find that when biosamples are added using only DNase-seq experiments, the resulting imputations for other assays can still be of high quality.

## 2 Methods

### 2.1 Avocado

#### Avocado topology

Avocado is a multi-scale deep tensor factorization model. The tensor factization component is comprised of five matrices of latent factors that encode the biosample, assay, and three resolutons of genomic factors at 25 bp, 250 bp, and 5 kbp resolution. Having multiple resolutions of genomic factors means that adjacent positions along the genome may share the same 250 bp and 5 kbp resolution factors. We used the same model architecture as in the original Avocado model [10], with 32 factors per biosample, 256 factors per assay, 25 factors per 25-bp genomic position, 40 factors per 250-bp genomic position, and 45 factors per 5-kbp genomic position. The neural network model has two hidden dense layers that each have 2,048 neurons, before the regression output, for a total of three weight matrices to be learned jointly with the matrices of latent factors. The network uses ReLU activation functions, ReLU(*x*) = max(0, *x*), on the hidden layers, but no activation function on the prediction.

#### Avocado training

Avocado is trained in a similar fashion to our previous work [10]. This procedure involves two steps, because the genome is large and the full set of genomic latent factors cannot fit in memory. The first is to jointly train all parameters of the model on the ENCODE Pilot regions, which comprise roughly 1% of the genome. After training is complete, the neural network weights, the assay factors, and the biosamples are all frozen. The second step is to train only the three matrices of latent factors that make up the genomic factors on each chromosome individually. In this manner, we can train comparable latent factors across each chromosome without the need to keep then all in memory at the same time.

Avocado was trained in a standard fashion for neural network optimization. All initial model parameters and optimizer hyperparameters were set to the defaults in Keras. In this work, Avocado was trained using the Adam optimizer [12] for 8,000 epochs with a batch size of 40,000. This is longer than our original work, where the model was trained for 800 epochs initially and 200 epochs on the subsequent transfer learning step. Empirical results suggest that this longer training process is required to reach convergence, potentially because of the large diversity of signals in the ENCODE2018-Core data set. When adding in additional biosample or assay factors, due to the small number of trainable parameters, the model was trained for only 10 epochs with a batch size of 512. Due to the large data set size, one epoch is defined as one pass over the genomic axis, randomly selecting experiments at each position, rather than one full pass over every experiment.

The model was implemented using Keras (https://keras.io) with the Theano backend [13], and experiments were run using GTX 1080 and GTX 2080 GPUs. For further background on neural network models, we recommend the comprehensive review by J. Schmidhuber [14].

### 2.2 Data and evaluation

#### ENCODE data set

We downloaded 6,870 genome-wide tracks of epigenomic data from the EN-CODE project (https://www.encodeproject.org). These experiments were all processed using the ENCODE processing pipeline and mapped to human genome assembly hg38, except for the ATAC-seq tracks, which were processed using an approach that would later be added to the ENCODE processing pipeline. The values are signal p-value for ChIP-seq data and ATAC-seq, read-depth normalized signal for DNase-seq, and plus/minus strand signal for RNA-seq. When multiple replicates were present, we preferentially chose the pooled replicate; otherwise, we chose the second replicate. The experimental signal tracks were then further processed before being used for model training. First, the signal was downsampled to 25 bp resolution by taking the average signal in each 25 bp bin. Second, a hyperbolic sin transformation was applied to the data. This transformation has been used previously to reduce the effect of outliers in epigenomic signal [9, 15].

We divided these experiments into two data sets, the ENCODE2018-Core data set and the ENCODE2018-Sparse data set. The ENCODE2018-Core data set contains 3,814 experiments from all 84 assays that have been performed in at least five biosamples, and all 400 biosamples that have been characterized by at least five assays. Hence, ~88.6% of the data in the ENCODE2018-Core data matrix is missing. The ENCODE2018-Sparse data set contains 3,056 experiments, including 1,281 assays that have been performed in fewer than five biosamples and 667 biosamples that have been characterized by fewer than five assays, yielding a matrix that is ~99.7% missing. These data sets, and the resulting model, can be found at https://noble.gs.washington.edu/proj/avocado/. The authors place no restrictions on the download and use of our generated data sets or model.

#### ENCODE-DREAM challenge data sets

For our comparisons with the ENCODE-DREAM challenge participants, we acquired from the challenge organizers both genome-wide model predictions from the top four participants and the binary labels (https://www.synapse.org/#!Synapse:syn17805945). The predictions and labels were defined at 200 bp resolution, with a stride of 50 bp, meaning that each 50 bp bin was included in four adjacent bins. The labels corresponded to conservative thresholded irreproducible discover rate (IDR) peaks called from multiple replicates of ChIP-seq signal.

#### Comparison to ENCODE-DREAM predictions

Avocado’s predictions had to be processed in several ways to make them comparable with the data format for the challenge. First, because Avocado’s predictions are in hg38 and the challenge was performed in hg19, the UCSC liftOver command (https://genome.ucsc.edu/cgi-bin/hgLiftOver) was used to convert the coordinates across reference genomes. Unfortunately, many of the 25 bp bins in hg38 mapped to the middle of bins in hg19, blurring the signal. Further, ~27% of positions on chromosome 21 of hg38 could not be mapped to positions in hg19, so those positions were discarded from the analysis. Lastly, because the challenge was performed at 200 bp resolution, the average prediction in the 200 bp region was used as Avocado’s predictions for that bin. We then filtered out all regions that were marked as “ambiguous”’ by the challenge organizers. These regions included both the flanks of true peaks as well as regions that were considered peaks in some, but not all, replicates.

The evaluation of each model was performed using both the average precision, which roughly corresponds to the area under a precision-recall curve, and the point along the precision-recall curve of equal precision and recall (EPR). The EPR corresponds to setting the decision threshold so that the number of positive predictions made by the model is equal to the number of positive labels in the data set. This is also called the “break-even point”. A strength of EPR, in comparison to taking the recall at a fixed precision, is that it accounts for the true sparsity in the label set. For example, if it is known beforehand that an experimental track generally has between 100 and 200 peaks across the entire genome, then a reasonable user may use the top 150 predictions from a model. However, if an experimental track had between 10,000 and 20,000 peaks, then a user may use the top 15,000 predicted peaks.

#### Calculation of average activity

In several of our experiments we compared model performance against the average activity of an assay. In all instances involving the ENCODE2018-Core data set, “average activity” refers to the average signal value at each locus across all biosamples in the training set for that particular experiment. Because the predictions across the entire ENCODE2018-Core data set are made using five-fold cross-validation, the training set differ for tracks from different folds. This approach ensures that the track being predicted is not included in the calculation of average activity which would make the baseline unfair. In instances involving the ENCODE2018-Sparse data set, “average activity” refers to the average activity across all tracks of that assay that were present in the entire ENCODE2018-Core data set.

## 3 Results

### 3.1 Avocado’s imputations are accurate and biosample specific

We first aimed to evaluate systematically the accuracy of Avocado’s imputed values on the ENCODE2018-Core data set. One challenge associated with this assessment is that no competing imputation method has yet been applied to this particular data set, making a direct comparison of methods difficult. However, we have shown recently that the average activity of a given assay across many biosamples is a good predictor of that activity in a new biosample [16]. Admittedly, this predictor is scientifically uninteresting, in the sense that it makes the same prediction for every new biosample and so, by construction, cannot capture biosample specific variation. We reasoned that improvement over this baseline indicates that the model must be capturing biosample specific signal. However, because the signal from most epigenomic assays is similar across biosamples, the average activity predictor serves as a strong baseline that any cross-cell type predictor must beat. Accordingly, we compare the predictions made by Avocado to the average activity of that assay in the training set that was used for model training.

Overall, we found that Avocado is able to impute signal accurately for a variety of different types of assays. We compared Avocado’s imputations to those of the average activity predictor across 37,249,359 genomic loci from chromosomes 12-22 using five-fold cross-validation among epigenomic experiments in the ENCODE2018-Core data set. Qualitatively, we observed strong visual concordance between observed and imputed values across a variety of assay types (Figure 2A). In particular, the imputations capture the shape of peaks in histone modification signal, such as those exhibited in H3K27ac and H3K4me3, the shape of peaks found in assays of transcription factors like ELF1 and CTCF, and exon-specific activity in gene transcription assays. As our primary quantitative measure, we compute the global mean-squared error (MSE) between the observed and imputed values. This value reduces from 0.0807 to 0.0653 (paired t-test p-value of 1e-157), a reduction of 19.1%, between the average activity predictor and Avocado (Figure 2B).

We also compute five complementary quantitative measures. Two measures emphasize the ability of an imputation method to correctly identify peaks in the data. One of these (mse1obs), defined as the MSE in the positions with the top 1% of observed signal, corresponds to a notion of recall. The complementary measure (mse1imp), defined as the MSE in position with the top 1% of imputed signal, corresponds to precision. Three additional measures focus on the MSE in regions of biological activity: the MSE in promoters (mseProm), gene bodies (mseGene), and enhancers (mseEnh). In aggregate, Avocado outperforms the average activity baseline on all six performance measures (p-values between 8e-65 for mse1imp and 1e-157 for mseGlobal) (Figure 2B/C).

When grouped by assay, we find that Avocado outperforms the average activity in 71 of the 84 experiments in our test set according to mseGlobal. Further investigation suggested that these problematic assays were mostly of transcription, indicating a weakness of the Avocado model, or assays that may have been of poor quality (Supplementary Note 1).

**Figure 2:**
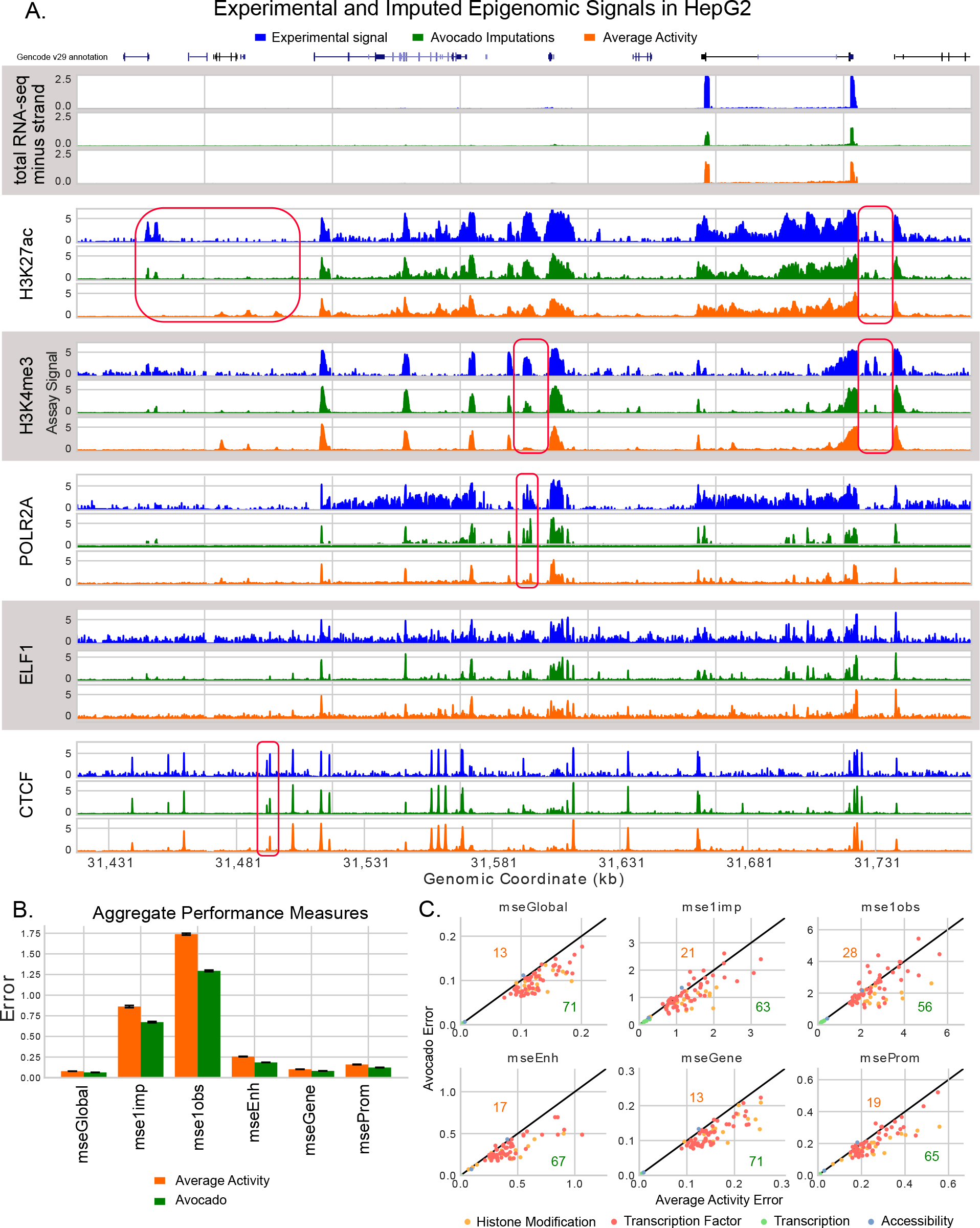
Avocado imputes epigenomic experiments accurately. (A) Example signal, corresponding imputations, and the average activity of that assay, for six assays performed in HepG2. The figure includes representative tracks for RNA-seq, histone modification, and factor binding. The data covers 350 kbp of chromosome 20. (B) Performance measures evaluated in aggregate over all experiments in chromosomes 12 through 22. Orange bars show the performance of the average activity baseline and green bars show the performance of Avocado’s imputations. (C) Performance measures evaluated for each assay, with Avocado’s error (y-axis) compared against the error of the average activity (x-axis). The number of assays in which Avocado outperforms the average activity is denoted in green for each metric, and the number of assays in which Avocado underperforms the average activity is denoted in orange.

The primary benefit of the ENCODE2018-Core data set, in comparison to previous data sets drawn from the Roadmap Compendium, is the inclusion of many more assays and biosamples. We hypothesized that not only will this data set allow us to make a more diverse set of imputations, but that these additional measurements will improve performance on assays already included in the Roadmap Compendium. We reasoned this may be the case because, for example, previous imputation approaches have imputed H3K36me3, a transcription associated mark, but have not utilized measurements of transcription to do so. A direct comparison to previous work was not simple due to differences in the processing pipelines and reference genomes, and so we re-trained Avocado using the same five-fold cross-validation strategy after having removed all experiments that did not originate from the Roadmap Epigenomics Consortium. Additionally, we removed all RNA-seq and methylation data sets, as they had not been used as input for previous imputation methods. This resulted in 1,072 tracks of histone modification and chromatin accessibility.

We found that the inclusion of additional assays and biosamples lead to a clear improvement in performance on the tracks from the Roadmap compendium. The MSE of Avocado’s imputations dropped from 0.115 when trained exclusively on Roadmap data sets to 0.107 when trained on all tracks in the ENCODE2018-Core data set, an improvement of 7% (p-value of 8e-45). When we grouped the error by assay, we observed that tracks appeared to range from a significant improvement to only a small decrease in performance (Supplementary Figure S2a). When aggregating these performances across assays, we similarly observe large improvements in the performance of most assays, and small decreases in a few (Supplementary Figure S2b/c). These results indicate that the inclusion of other phenomena do, indeed, aid in the imputation of the original tracks.

### 3.2 Comparison to ENCODE-DREAM participants

Predicting the binding of various transcription factors is particularly important due both to these proteins’ critical roles in regulating gene expression and the sparsity with which their binding has been experimentally characterized across different biosamples. For example, of the 50 transcription factors included in the ENCODE2018-Core data set, only 11 have been performed in more than 10 biosamples. The most performed assay measures CTCF binding, and has been performed 136 times, which is almost twice as high as the next most performed assay, measuring POLR2A binding, at 70 assays. In contrast, 11 of the 18 histone modifications in the ENCODE2018-Core data set have been measured in more than 10 biosamples, and the top 6 have all been performed in more than 200 biosamples. The sparsity of protein binding assays is exacerbated in the ENCODE2018-Sparse data set, where an additional 707 assays measuring protein binding have been performed in fewer than five biosamples.

A recent ENCODE-DREAM challenge focused on the prediction of transcription factor binding across biosamples, and phrased the prediction task as one of classification where the aim is to predict whether binding is occuring at a given locus (https://www.synapse.org/#!Synapse:syn6131484). The challenge involved training machine learning models to predict signal peaks using nucleotide sequence, sequence properties, and measurements of gene expression and chromatin accessibility. The participants trained their models on a subset of chromosomes and biosamples, and were evaluated based on how well their models generalized both across chromosomes and in new biosamples. We acquired predicted probabilities of binding from the top four teams, Yuanfang Guan, dxquang, autosome.ru, and J-TEAM, for 13 tracks of epigenomic data. Avocado did not make predictions for four of the included assays, E2F1, HNF4A, FOXA2, NANOG, so only 9 tracks were used in this evaluation.

We compared Avocado’s predictions of transcription factor binding to the predictions of the top four models from the ENCODE-DREAM challenge to serve as an independent validation of Avocado’s quality. We used both the average precision (AP) and the point on the precision-recall curve where precision and recall are equal (EPR) to evaluate the methods. In order to provide an upper limit for how good Avocado’s predictions could be after the conversion process, we included as a baseline the experimental ChIP-seq data that the peaks were called from (called “Same Biosample”). Additionally, we compared against the average activity of that assay in Avocado’s training set for that prediction. This baseline serves to show that Avocado is learning to make biosample-specific predictions. Further, when we investigated the training sets for the various experiments, we noted that there were two liver biosamples, male adult (age 32), and female child (age 4), that had similar assays performed in them. To ensure that Avocado was not simply memorizing the signal from one of these biosamples and predicting it for the other liver biosample, we compare against the signal from the related biosample as well (denoted “Similar Biosample”).

We observed that Avocado’s predictions outperform all of the challenge participants in all tracks except for CTCF in iPSC and FOXA1 in liver (Table 1, Supplementary Table S1). The most significant improvement comes in predicting REST, a transcriptional factor that represses neuronal genes in biosamples that are not neurons, and the highest overall performance is in predicting CTCF binding, likely because CTCF binding is similar across most biosamples. Importantly, the REST assay for both liver biosamples were in the same fold, and TAF1 was only performed in one of the liver biosamples, so Avocado’s good performance on those tracks are strong indicators of its performance. Visually, we observe that some of the participants models appeared to over-predict signal values, suggesting that a source of error for these models is their lack of precision, corresponding to rapid drop in precision for predicting REST (Supplementary Figure S3). Interestingly, Avocado appears to underperform using the related liver biosample as the predictor for FOXA1, suggesting that perhaps the factors for FOXA1 are poorly trained. However, this result is further evidence that Avocado is not simply memorizing related signal. We also note that, in the case of CTCF in iPSCs, the ChIP-seq signal from iPSC appears to underperform two challenge participants, suggesting that the conversion process may limit Avocado’s performance.

**Table 1:** Comparison of methods on ENCODE-DREAM challenge test set. The average precision (AP) computed across nine epigenomic experiments in the ENCODE-DREAM challenge test set in chromosome 21. For each track, the score for the best-performing predictive model is in boldface.

| Biosample<br>Assay<br>Method | iPSC<br>CTCF | PC-3<br>CTCF | liver<br>EGR1 | liver<br>FOXA1 | liver<br>GABPA | liver<br>JUND | liver<br>MAX | liver<br>REST | liver<br>TAF1 |
| --- | --- | --- | --- | --- | --- | --- | --- | --- | --- |
| Yuanfang Guan | 0.729 | 0.600 | 0.397 | 0.282 | 0.353 | 0.533 | 0.441 | 0.319 | 0.281 |
| dxquang | 0.866 | 0.783 | 0.274 | 0.400 | 0.347 | 0.260 | 0.330 | 0.312 | 0.264 |
| autosome.ru | 0.778 | 0.486 | 0.331 | 0.243 | 0.342 | 0.416 | 0.384 | 0.264 | 0.221 |
| J-TEAM | **0.812** | 0.747 | 0.363 | **0.462** | 0.344 | 0.415 | 0.377 | 0.196 | 0.272 |
| Avocado | 0.723 | **0.791** | **0.530** | 0.354 | **0.396** | **0.660** | **0.574** | **0.477** | **0.384** |
| Similar Biosample | — | — | 0.363 | 0.389 | 0.226 | 0.568 | 0.446 | 0.408 | — |
| Same Biosample | 0.741 | 0.878 | 0.648 | 0.716 | 0.573 | 0.731 | 0.622 | 0.622 | 0.556 |
| Average Activity | 0.574 | 0.735 | 0.240 | 0.299 | 0.253 | 0.223 | 0.349 | 0.124 | 0.140 |

We did our best to ensure a fair comparison between Avocado and the challenge participants, but the comparison is necessarily imperfect, for several reasons. Two factors make the comparison easier for Avocado. First, Avocado is exposed to many epigenomic measurements that the challenge participants did not have available, including measurements of the same transcription factor in other cell types. Second, as an imputation approach, Avocado is trained on the same genomic loci that it makes predictions for, whereas the challenge participants had to make predictions for held-out chromosomes. On the other hand, three factors skew the comparison in favor of the challenge participants. First, unlike the challenge participants, Avocado was not directly exposed to any aspect of nucleotide sequence or motif presence. Second, Avocado makes predictions at 25 bp resolution in hg38, whereas the challenge was conducted at 200 bp resolution in hg19. We were able to use liftOver to convert between assemblies, followed by aggregating the signal from 25 bp resolution to 200 bp resolution, but both steps blurred the signal. Third, Avocado is trained to predict signal values directly, whereas the challenge participants are trained on the classification task of identifying whether a position is a peak. Evaluation is done in a classification setting. In particular, Avocado is penalized for accurately predicting high signal values in regions that aren’t labeled as peaks, exemplifying the discordance between the regression and classification settings. For all these reasons, Avocado would not have been a valid submission to the challenge. Finally, it is perhaps worth emphasizing that whereas the challenge was truly blind, our application of Avocado to the challenge data is only blind “by construction.” We emphasize that we did not adjust Avocado’s model or hyperparameters based on looking at the challenge results: the comparison presented here is based entirely on a pre-trained Avocado model.

### 3.3 Extending Avocado to more biosamples and assays

Despite including 3,814 epigenomic experiments, the ENCODE2018-Core data set does not contain all biosamples or assays that are represented in the ENCODE compendium. Specifically, the data set does not include 667 biosamples where fewer than five assays had been performed, and it does not include 1,281 assays that had been performed in fewer than five biosamples. The missing biosamples primarily include time courses, genetic modifications, and treatments of canonical biosamples, such as HepG2 genetically modified using RNAi. However, several primary cell lines and tissues such as amniotic stem cells, adipocytes, and pulmonary artery, were also not included in the ENCODE2018-Core data set due to lack of sufficient data. The majority of the missing assays corresponded to transcription measurements after gene knockdowns/knockouts (shRNA and CRISPR assays) or to binding measurements of eGFP fusion proteins. Yet some transcription factors, such as NANOG, FOXA2, and HNF4A, were excluded as well. We collect these experiments into a separate data set, called ENCODE2018-Sparse (see Methods 2.2).

We constructed the ENCODE2018-Sparse data set to attempt to address some of the problems of missingness in ENCODE2018-Core. This sparse version of the data has 99.7% missing entries, in comparson to 88.6% missing in ENCODE2018-Core. Within ENCODE2018-Sparse, we identified four main groups of biosamples: (1) 417 biosamples that only had DNase-seq performed on them, with 58 additional biosamples that had DNase and one or more other assays performed in it (2) 112 biosamples that had various measurements of transcription performed in them, (3) 7 biosamples that were well characterized by at least 50 sparsely performed assays of transcription factor binding, and (4) biosamples derived from HepG2 and K562 that were well characterized by various knockouts (Supplementary Figure S1).

In general, handling sparsely characterized assays or biosamples in a model like Avocado is challenging. Hence, we designed a three-step process that we hypothesized would allow us to make accurate imputations for additions with few corresponding tracks (Supplementary Figure S4). This approach is conceptually similar to our main approach for training Avocado. First, we trained the Avocado model on all 3,814 experiments in ENCODE2018-Core. Second, we froze all of the weights in the model, including both the neural network weights and all five of the latent factor matrices. Third, we fit the new biosample or assay factors to the model using only the experimental signal derived from the ENCODE Pilot Regions. This resulted in a model whose only difference was the inclusion of a set of trained assay or biosample factors that were not present in original model. This training strategy has the benefit of allowing for quick addition of biosamples or assays to the pre-trained model, with requiring retraining of any of the existing model parameters.

In order to test the effectiveness of this approach, we extended Avocado to include assays that were in the ENCODE-DREAM challenge but not in the ENCODE2018-Core data set. For the four assays that we did not compare against (HNF4A and FOXA2 in liver, NANOG in iPSC, and E2F1 in K562), all but E2F1 had been performed in a biosample other than the one included in the challenge. Fortunately, for the two tracks derived from liver, the assays had also been performed in a non-liver biosample, allowing us to more rigorously test Avocado’s generalization ability. Accordingly, we fit these three new assay factors using the procedure above. This fitting was done using HNF4A and FOXA2 from HepG2 and NANOG from h1-hESC. We then used the new assay factors, coupled with the pre-trained network, genome factors, and relevant biosample factors, to impute three remaining tracks in the challenge.

We observed that Avocado’s imputed tracks for HNF4A and FOXA2 in liver were of high quality and outperformed several baselines (Figure 3a). Most notably, both of these tracks outperformed all four challenge participants in their respective settings according to both EPR and AP. Second, they both outperformed simply using the track that they were trained on as the predictor, indicating that the model is leveraging the pre-trained biosample latent factors to predict biosample specific signal. Due to FOXA2’s similarity to FOXA1, we anticipated that predictions for these two transcription factors would be similar. To ensure that the model was not simply predicting the same signal for both FOXA1 and FOXA2, we compared against the predictions made for FOXA1 and noted that they perform worse than the predictions for FOXA2. Interestingly, the predictions for FOXA1 appear to outperform all four challenge participants’ predictions for FOXA2, highlighting that the two transcription factors have very similar binding propensities.

However, we also observed that Avocado’s imputations for NANOG in iPSCs are of particularly poor quality. Avocado’s predictions underperform all four challenge participants. More notably, Avocado also underperforms using the signal from h1-hESC that it was trained on as the predictor. One potential reason for this poor performance is that relevant features of the NANOG binding sites are not encoded in the genomic latent factors. Alternatively, given that Avocado also underperformed the challenge participants at predicting CTCF in iPSC, it may be that the iPSC latent factors are not well trained, leading to poor performance in predictions of any track.

Finally, we tested the ability of the three-step process in Supplementary Figure S4 to make accurate predictions for biosamples that the model was not originally trained on. We observed good performance when imputing assays for biosamples that were trained using only DNase-seq (Supplementary Note 2).

## 4 Discussion

To our knowledge, we report here the largest imputation of epigenomic data that has been performed to date. We applied the Avocado deep tensor factorization model to 3,814 epigenomic experiments in the ENCODE2018-Core data set. The resulting imputations cover a diverse set of biological activity and cellular contexts. Due to the cost of experimentation and the increasing sparsity of epigenomic compendia we anticipate that imputations of this scale will serve as a valuable communal resource for characterizing the human epigenome.

**Figure 3:**
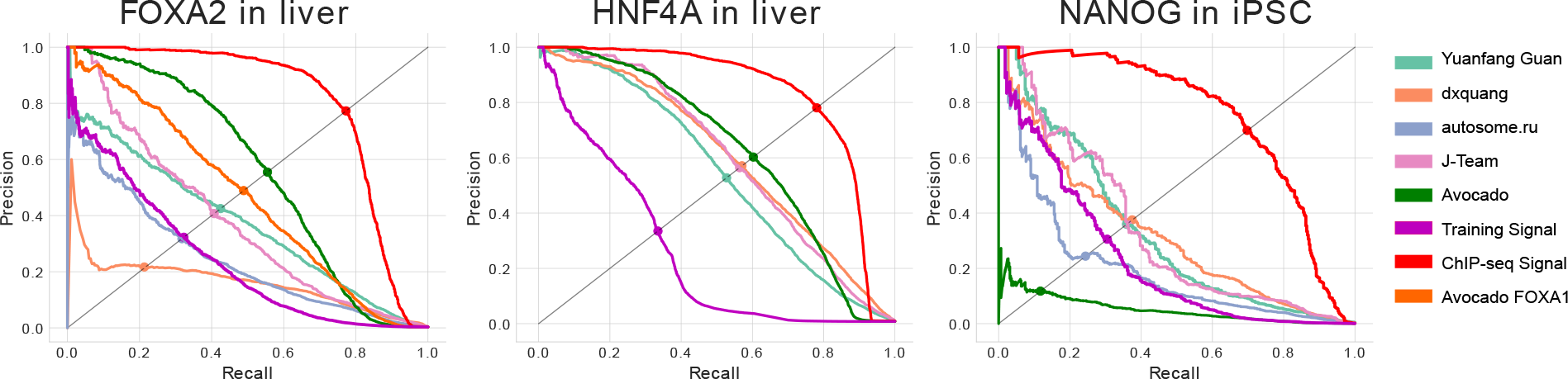
Avocado’s performance when adding new transcription factors to a pre-trained model. Precision-recall curves for three transcription factors that were added to a pre-trained model using a single track of data each from the ENCODE2018-Sparse data set.

We used multiple independent lines of reasoning to confirm that Avocado’s imputations are both accurate and biosample specific. First, we compared each imputed data track to the average activity of that assay and found that, for almost all assays, that Avocado’s imputations were more accurate. A current weakness in Avocado’s imputations is imputing transcription, likely due to the sparse, exon-level activity of these assays along the genome. Second, we compared imputations of transcription factor binding tracks to the predictions made by the top four models in the recent ENCODE-DREAM challenge. In almost all cases, the Avocado imputations were significantly more accurate than the imputations produced by the challenge participants. Notably, Avocado is not exposed to nucleotide sequence at all during the training process, and so its ability to correctly impute transcription factor binding is based entirely on local epigenomic context, rather than binding motifs.

Ongoing characterization efforts regularly identify new biosamples of interest and develop assays to measure previously uncharacterized phenomena. These efforts aid in understanding the complexities of the human genome but pose a problem for imputation efforts that must be trained in a batch fashion. Given that it took almost a day to fit the Avocado genomic latent factors for even the smallest chromosome, re-training the model for each inclusion is not feasible. We demonstrated that, by leveraging parameters that had been pre-trained on the ENCODE2018-Core data set, new assays and biosamples could be quickly added to the existing Avocado model. Our observations suggest that not only is this approach computationally efficient, with three new assays taking only a few minutes to add to the model, but that the resulting imputations are highly accurate.

One potential reason that this pre-training strategy works well is that the genomic latent factors efficiently encode information about regions of biological activity. For example, rather than memorizing the specific assays that exhibit activity at each locus, the latent factors may be organizing general features of the biochemical activity at that locus. We have previously demonstrated the utility of Avocado’s latent genomic representation for several predictive tasks [10]. Investigating the utility and meaning of the latent factors from this improved Avocado model is ongoing work.

Notably, however, the encoding of relevant information in the latent factors may lead to a potential weakness in Avocado’s ability to generalize to novel biosamples or assays. Specifically, if the signal in a novel biosample or assay is not predictable from the tracks that were used to train the initial genomic latent factors, then it is unlikely that Avocado will make good imputations for the new data. For example, if a transcription factor is dissimilar to any factors in the training set, then the genomic latent factors may not have captured features relevant to the novel factor. This may explain why Avocado fails to generalize well to NANOG.

A strength of large consortia, such as ENCODE, is that they are able to collect massive amounts of experimental data. This amount of data is only possible because many labs collect it over the course of several years. Inevitably, this results in some data that is of poor quality. While quality control measures can usually identify data that is of very poor quality, they are not perfect, and the decision of what to do with such data can be challenging. Unfortunately, data of poor quality poses a dual challenge for any large scale imputation approach. When an imputation approach is trained on low quaity data, then the resulting imputations may be distorted by the noise. Furthermore, when the approach is evaluated against data that is of poor quality, imputations that are of good quality may be incorrectly scored poorly. Thus, when dealing with large and historic data sources, it is important to ensure the quality of the data being used.

The imputation approach offered by Avocado has great potential to be extended to precision medicine. In this setting, a biosample is sparsely assayed in a variety of individuals, and the goal is to correctly impute the inter-individual variation, particularly in regions associated with disease. We anticipate that Avocado could either be applied directly in this setting, or be extended to accommodate a 4D data tensor, where the fourth dimension corresponds to distinct individuals.

## Acknowledgments

This work was supported in part by an NSF IGERT grant (DGE-1258485) and by NIH awards U54 DK107979 and U41 HG007000. We would also like to thank Giancarlo Bonora, Timothy Durham, Ritambhara Singh, and Gürkan Yardımcı for many productive discussions, as well as Anshul Kundaje and Akshay Balsubramani for providing data from the ENCODE-DREAM challenge.

## Supplementary Note 1

We investigated further the 13 experiments for which Avocado underperforms the average activity predictor. This set is enriched for measurements of transcription: 10 of the 13 experiments (77%) measure gene transcription, such as CAGE, RAMPAGE, microRNA-seq, polyA-depleted RNAseq, and small RNA-seq. The remaining three assays for which Avocado does not outperform the average activity predictor according to mseGlobal are H3K9me2, EP300, and ATAC-seq. Further investigation on the ENCODE portal showed all H3K9me2 experiments had audit warnings and that only one of the experiments, in iPSC cells, had a fraction of reads in peaks (FRiP) score above the general quality control theshold of 1% used for ChIP-seq experiments [17]. While standards for ATAC-seq experiments have been released, the quality metrics associated with the experiments we used had not yet been released on the ENCODE portal, and so we were unable to verify their quality.

We then investigated those assays that Avocado underperformed the average activity baseline on other performance measures. First, we notice that Avocado imputed transcription poorly across all measures. On all measures except mse1imp, at least 9 of the underperforming assays related to measurements of transcription. Second, we notice that H3K9me2 and ATAC-seq are poor performers across all metrics as well. The consistent poor performance of these 11 assays may give a more pessimistic view of Avocado’s performance in general.

We then evaluated assay performance across different performance measures. We noticed that Avocado only underperforms the average activity baseline on only five of the problematic transcription assays. This suggests that Avocado may have a higher precision than recall when it comes to predicting exon-specific activity. However, one weakness of mse1imp and mse1obs is that the percentage used to approximate peak coverage, 1%, may be appropriate for histone marks, but is not as specific to areas of transcription.

### Supplementary Note 2

We wanted to evaluate the ability for the three step processes descrived in Supplemantary Figure S4 to fit new biosamples from a limited number of assay measurements. To do so, we began by training biosample factors for 475 biosamples not in the ENCODE2018-Core data set that had DNase-seq performed in them. We then evaluated Avvocado’s ability to predict other assays that were performed in these biosamples. A large number of these biosamples had only DNase-seq performed in them, so we also evaluated Avocado’s ability to predict DNase-seq as well. We reasoned that because the biosample factors were trained using the ENCODE Pilot regions, but the predictions were evaluated in chromosome 20 without re-training the corresponding genomic latent factors, this would be a fair evaluation.

We observed good performance of the imputations for these biosamples. Visually, we notice the same concordance between the imputed and the experimental signal and that biosample-specific elements are being captured (Supplementary Figure S5a). We then evaluated the performance of Avocado on the mseGlobal metric compared to the average activity baseline for each assay. We observed that Avocado appears to produce high quality predictions for several assays, including CTCF, H3K27ac, and POLR2A (Supplementary Figure S5b). However, for other assays, such as H3K9me3 an H3K36me3, the average activity dominates. It is possible that this phenomenon speaks to the ability of DNase to recover these other approaches. Overall, we observe a decrease in error from 0.027 when using the average activity to 0.024 (paired t-test p-value of 1e-5) when using the imputations from Avocado.

**Table S1:**
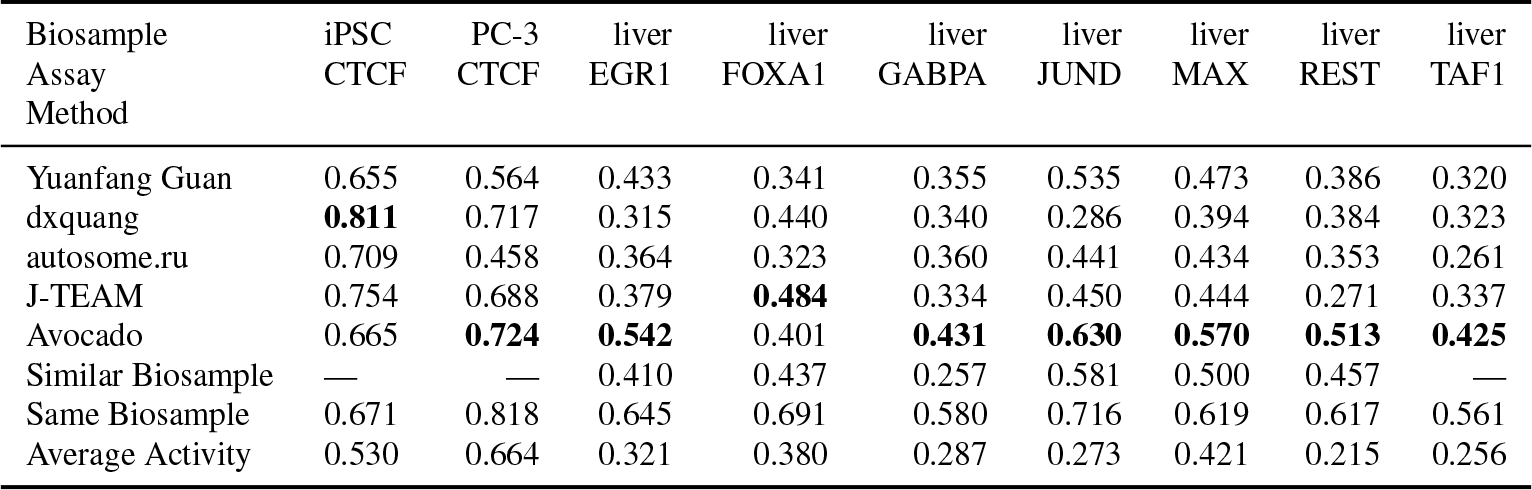
Comparison of methods on ENCODE-DREAM challenge test set. The equal precision-recall (EPR) computed across nine epigenomic experiments in the ENCODE-DREAM challenge test set in chromosome 21. For each track, the score for the best-performing predictive model is in boldface.

**Figure S1:**
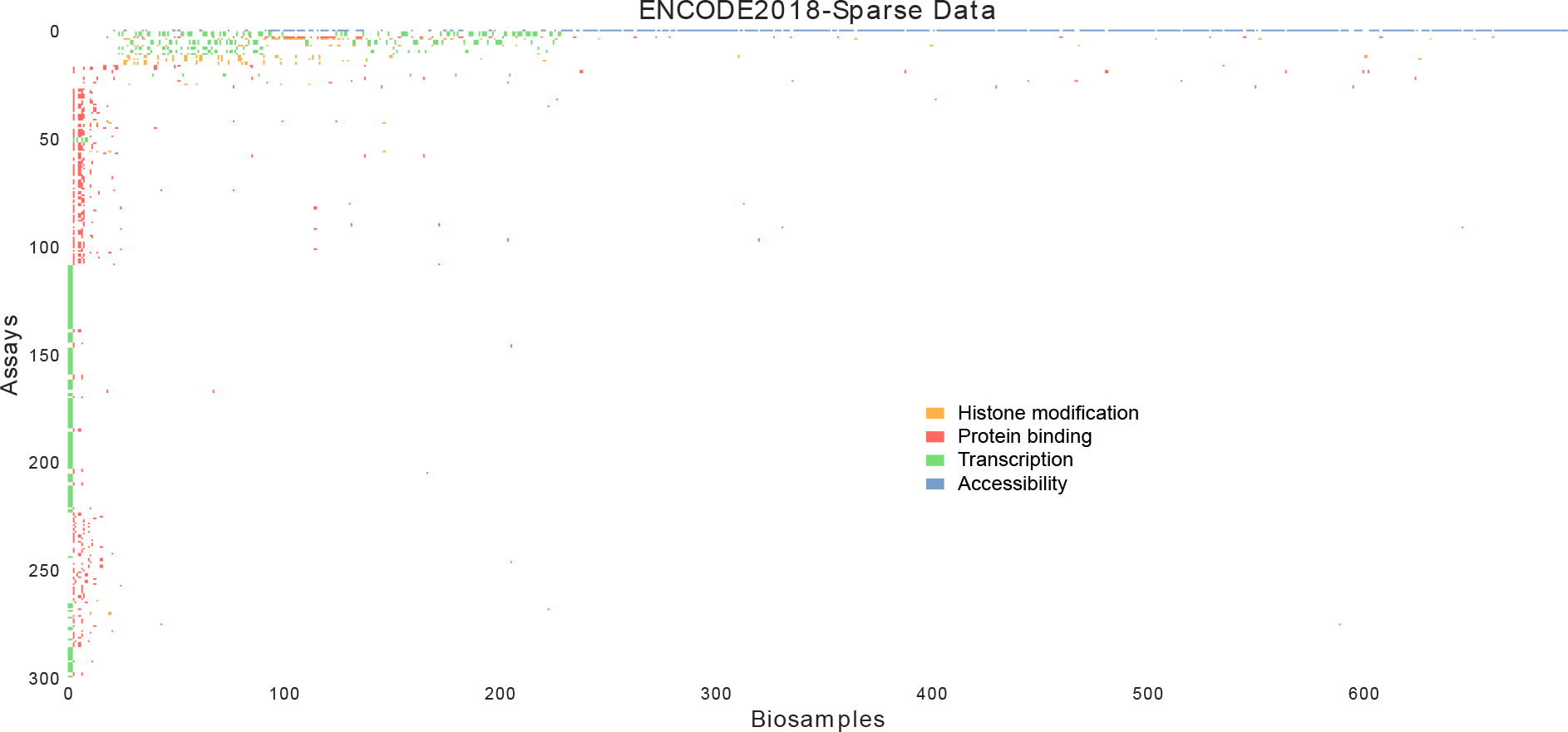
The ENCODE2018-Sparse data matrix. The ENCODE2018-Sparse data matrix includes all assays that were performed in fewer than 5 biosamples, and all biosamples that were characterized by fewer than 5 assays. Experiments that have been performed are displayed as colored rectangles, and experiments that have not been performed are displayed as white. The color corresponds to the general type of assay, with blue indicating chromatin accessibility, orange indicating histone modification, red indicating protein binding, and green indicating transcription. This figure displays all biosamples, and the top 300 assays ranked number of biosamples that they were performed in.

**Figure S2:**
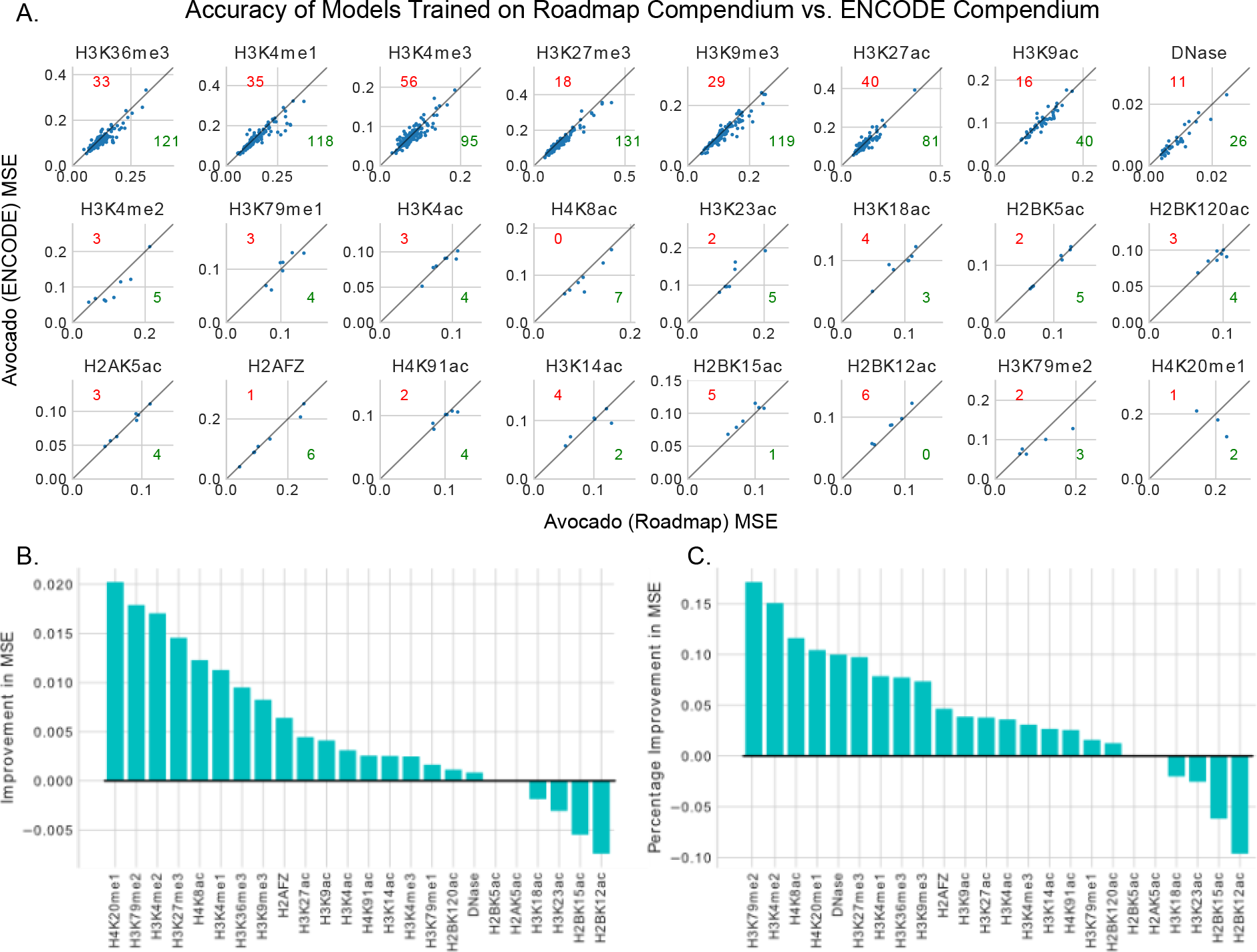
Accuracies of models trained on either the Roadmap compendium or the ENCODE2018-Core data. (A) Each panel depicts the error of models trained on either the ENCODE2018-Core data set (Avocado (ENCODE)), or those tracks from the ENCODE2018-Core data set that were provided by the Roadmap Epigenomics Consortium (Avocado (Roadmap)), when imputing the tracks contained in the latter. Each dot corresponds to MSE on a single track, and each panel corrsponds to all tracks from that assay. Dots below the diagonal line indicate that the model trained on the ENCODE2018-Core data set outperformed the model trained on the Roadmap data set, with the number in green specifying the number of such tracks, and dots above the line indicate the reverse, specified by the red number. (B) The improvement in performance when using a model trained on the full ENCODE2018-Core data set versus one trained on only the Roadmap tracks. (C) Similar to (B), except the percentage improvement.

**Figure S3:**
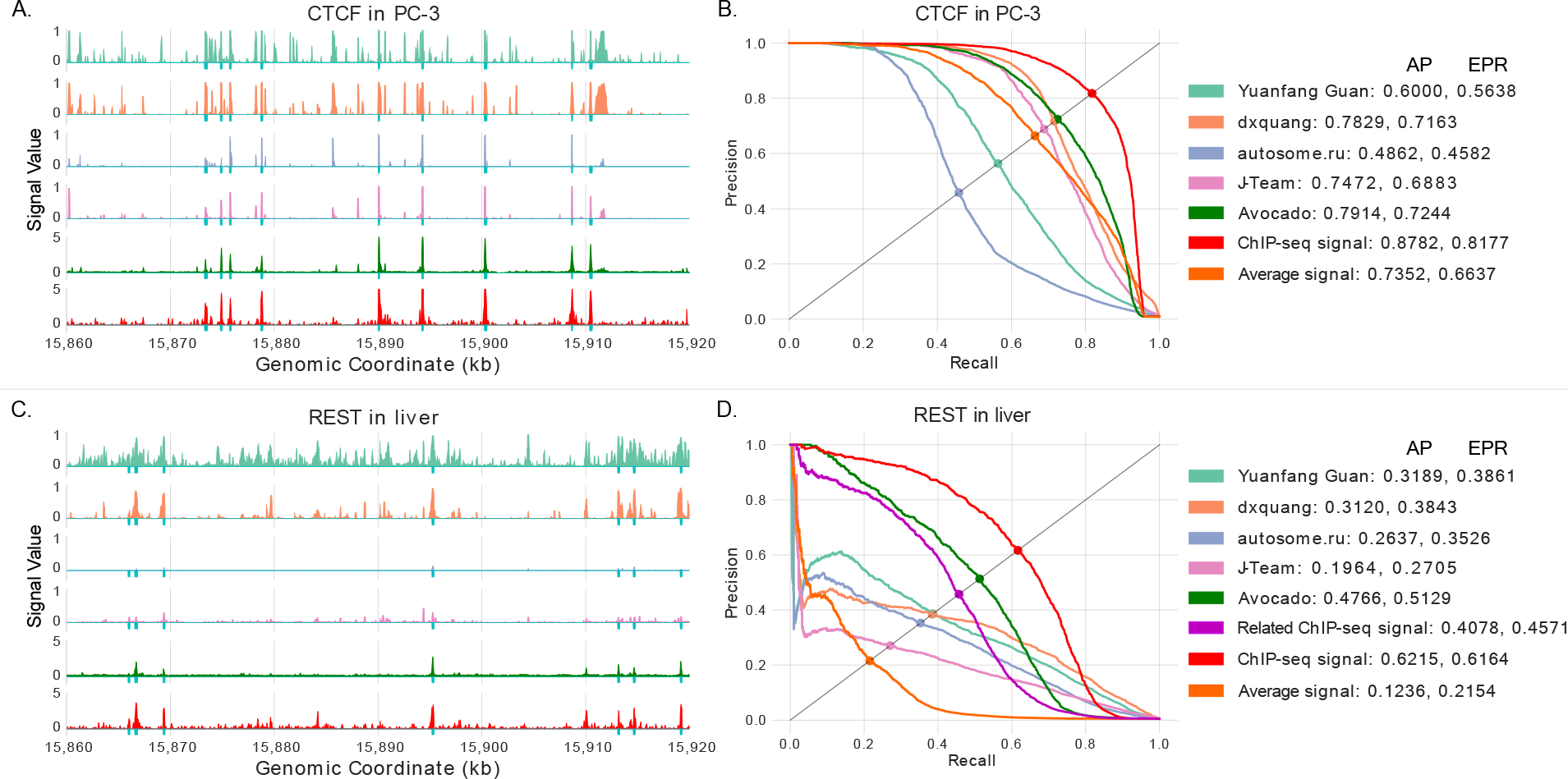
Avocado imputes transcription factors correctly. (A) Example predictions from a region of chromosome 21 for the top four ENCODEDREAM participants, Avocado, and experimental ChIP-seq data measuring CTCF binding in PC-3. Cyan ticks at the bottom of the tracks indicate peak calls. (B) A precision-recall curve showing the performance of the four participants and Avocado in chromosome 21. As additional baselines, the experimental ChIP-seq signal (red) and the average signal across Avocado’s training set (orange) were included in the comparison. For each approach, the average precision (AP) and the equal-precision-recall (EPR) are reported (see Section 2.2), and the position on the curve where the EPR lies is marked as a dot. (C) Similar to (A), except for REST binding in a liver biosample. (D) Similar to (B), except for REST binding in a liver biosample. The experimental signal from a different liver biosample is used as a further baseline (magenta).

**Figure S4:**
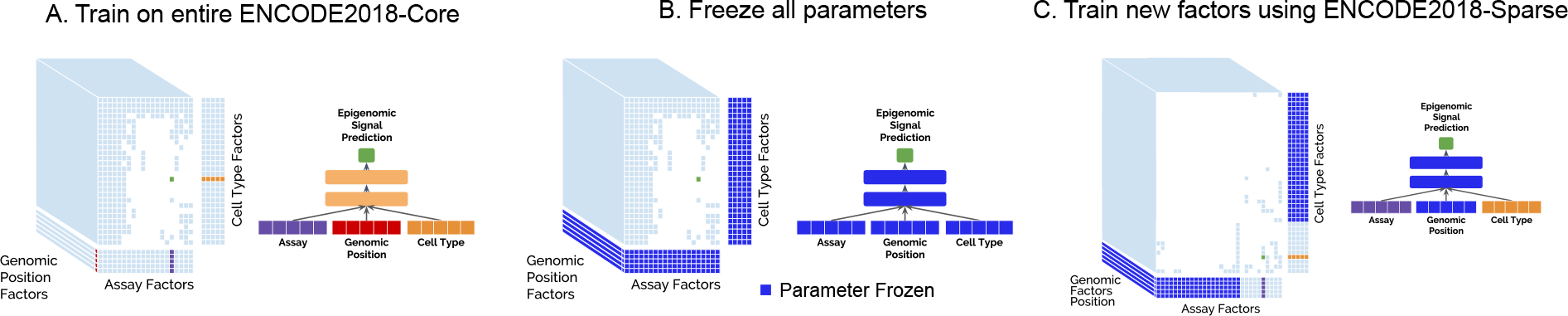
Transfer learning methodology. >A schematic of the three step process to train Avocado on the ENCODE2018-Sparse data set. (A) Train Avocado on the entire ENCODE2018-Core data set as normal. (B) Freeze the weights of both the neural network and the factors. (C) Train only the factor values for new biosamples and assays that are being added to the model.

**Figure S5:**
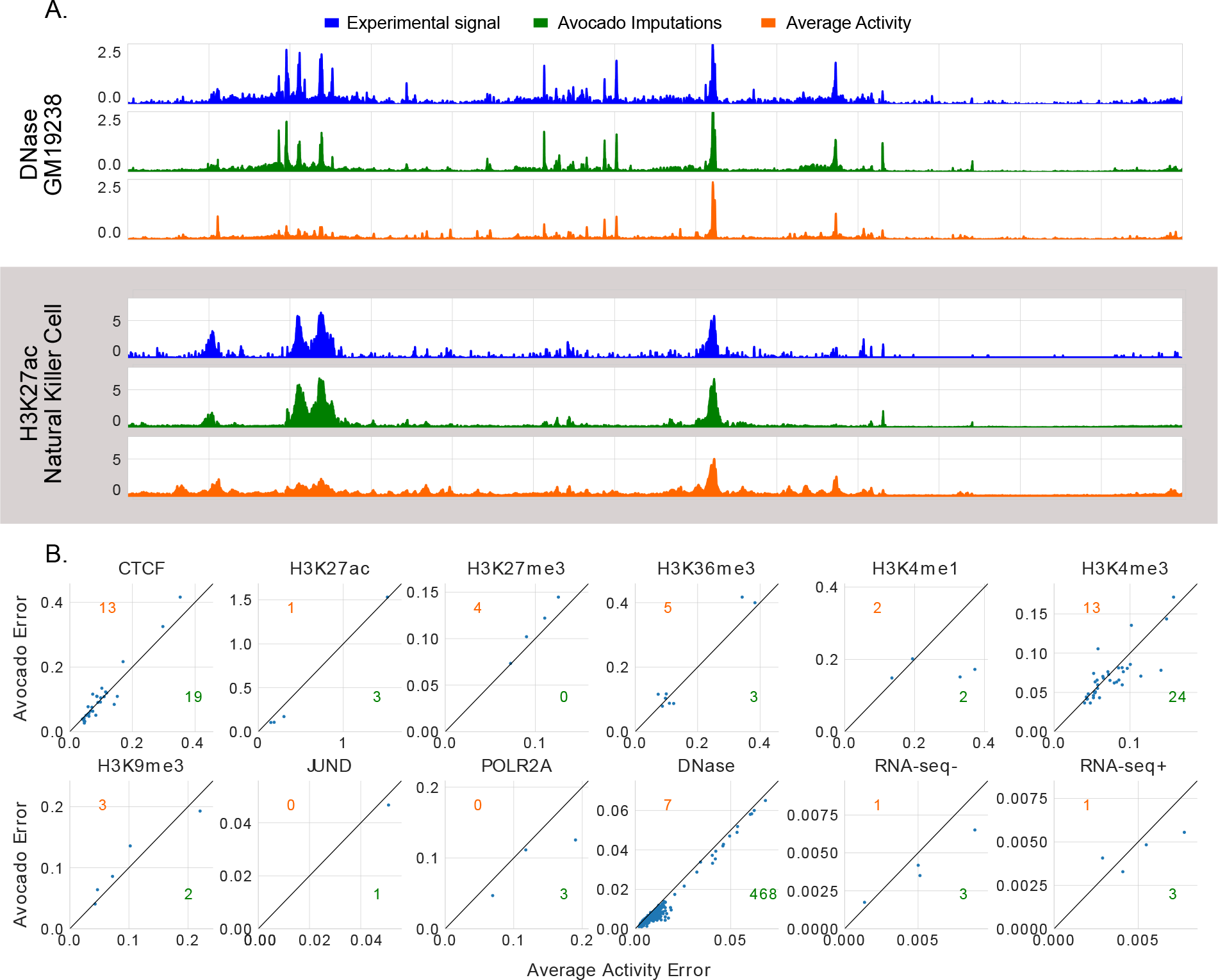
Imputations and performance when adding biosamples to a pre-trained model. (A) Imputations for two tracks of data in the ENCODE2018-Sparse data set on chromosome 20 after fitting the biosample factors using only DNase-seq signal from the ENCODE Pilot Regions. (B) Performance of Avocado at imputing tracks on chromosome 20 after fitting the biosample factors using only DNase-seq signal from the ENCODE Pilot regions.

